# Differential activation of p53-Lamin A/C and p16-RB mediated senescence pathways in trophoblast from pregnancies complicated by type A2 Gestational Diabetes Mellitus

**DOI:** 10.1101/2025.03.04.641461

**Authors:** Leena Kadam, Kaylee Chan, Kylia Ahuna, Nicole Marshall, Leslie Myatt

## Abstract

Gestational diabetes mellitus (GDM) increases maternal risks such as hypertension and future type 2 diabetes while also contributing to fetal complications such as large-for-gestational-age infants and stillbirth. The placenta which is crucial for fetal development, exhibits structural and functional changes in GDM, but the impact of these alterations on placental trophoblast function remains unclear. During their differentiation villous cytotrophoblast display several characteristics of senescent cells however the role of senescence pathways in placental function remains unexplored in GDM. Here we investigate whether placental senescence pathways are altered in GDM, utilizing term villous tissue and primary trophoblasts to assess molecular changes, and determined fetal sex-based differences. Our data suggest that both p21 and p16 mediated senescence pathways are activated during trophoblast differentiation and are dysregulated in GDM placenta in a sexually dimorphic manner. We also provide evidence for increased activation of p53-Lamin A/C and p16-RB pathways in trophoblast from GDM placentas. Reduced expression of p21 and its downstream effects on GCM1 expression and βhCG secretion outline how altered physiological senescence can affect trophoblast differentiation and function. This is a seminal study highlighting how placental senescence pathways are altered in pregnancies complicated by GDM.

## Introduction

Gestational diabetes mellitus (GDM) is a metabolic disorder of pregnancy which manifests as impaired glucose tolerance with onset or first recognition during pregnancy [1]. It is considered as one of the great obstetrical syndromes with an increasing incidence, currently being ∼14.7% worldwide and ∼7% of pregnancies in the USA [2,3]. Women with GDM are more susceptible to hypertensive disorders, dysfunctional labor and thromboembolic events during pregnancy and are also at higher risk of developing type 2 diabetes and cardiovascular disease in later life [4–7]. GDM can lead to fetal complications such as large-for-gestational-age (LGA) infants (which can lead to cesarean delivery, birth trauma, shoulder dystocia and asphyxia), fetal growth restriction (FGR) or stillbirth [4,5,8]. The offspring of women with GDM are also at a higher risk for development of obesity and diabetes in later life [4,5,8–10]. It has been observed that women carrying male fetuses are at a higher risk of developing GDM and associated co-morbidities highlighting that fetal sex might influence the development and severity of the disease [11,12]. The placenta, being the maternal-fetal interface, ensures optimal fetal nutrition, growth and development. The maternal metabolic conditions of hyperglycemia and hyperinsulinemia in GDM appear to affect placental development as GDM placentas are heavier and show histopathological defects such as immature villi, villous fibrinoid necrosis, increased angiogenesis and chorangiosis [13,14]. Extensive reduction of microvilli on thin hyper-vacuolized syncytiotrophoblast, a thick cytotrophoblast layer and a thick trophoblast basal membrane have also been reported in GDM placentas [15,16]. However, the effect of these changes on placental trophoblast function at the cellular level is not completely understood.

Cellular senescence is an irreversible cycle arrest wherein the cells stop proliferating but continue to be metabolically active. It is a physiological process integral to normal tissue homeostasis and functions with crucial roles in tumor suppression, wound healing and embryonic development [17–19]. Disruption in physiological senescence pathways is associated with declining tissue function in diseases such as hypertension and Type 2 Diabetes Mellitus in non-pregnant individuals [20,21]. Furthermore, stressors such as increased oxidative stress, inflammation, metabolic imbalance and insulin resistance - characteristic of pregnancies with GDM, are also known to increase senescence in non-pregnant individuals but their impact on placental senescence remains relatively unexplored in GDM tissues [22–30]. Interestingly, placental syncytiotrophoblast display several characteristics of senescent cells in being multinucleate, terminally differentiated cells with irreversible replication arrest. In recent years, elevated senescence has been reported in placentae from pathologies such as FGR, stillbirth, pre-eclampsia and GDM [31–33]. However, the role of senescence in placental function has not been evaluated.

In the present study, we aimed to determine if placental senescence pathways are altered in GDM potentially affecting its function. We utilized villous tissue as well as primary trophoblast isolated at term from placentas of normoglycemic and GDM pregnancies to evaluate changes in senescence pathway mediators. As it is known that pregnancies with male fetuses are more susceptible to developing GDM, we also evaluated fetal sex-based effects on trophoblast senescence.

## METHODS

### Patient recruitment and collection of placental tissue

This study was performed in line with the principles of the Declaration of Helsinki. Approval was granted by the Institutional Review Board of Oregon Health & Science University (IRB 16328). Informed consent was obtained from all patients before tissue collection. Placentas (n=10 per group with n=5 per sex) were collected from normoglycemic women (NW) and women with type A2 GDM (GDM). GDM was defined using IADPSG criteria which recommends 75-grams two hours OGTT with at least one abnormal result: Fasting plasma glucose (FPG) at ≥ 92 mg/dl or 1-hour ≥ 180 mg/dl or 2-hour ≥ 153mg/dl to be classified as GDM [34]. Women with type A2 GDM were those with greater than 30% of FPG ≥90-95 mg/dl or 1 hr Postprandial ≥130-140 mg/dl and needed medication to control their blood glucose levels. To keep the samples consistent, we focused on women with type A2GDM and collected samples from women who received insulin as medication to control their diabetes. Patients from both NW and GDM groups satisfied the following inclusion/exclusion criteria.

<u>Inclusion criteria</u> included a singleton pregnancy, an age range of 18-45 and delivery by cesarean section at term with no labor. <u>Exclusion criteria</u> included concurrent disease (including, but not limited to IUGR, hypertension, pre-eclampsia, eating disorders, infection, inflammatory disorders), use of tobacco, drugs, or medications other than to treat GDM, excessive weight gain or loss prior to pregnancy (>20 lbs), or bariatric surgery in the last year and labor defined by regular uterine contractions (every 3-4 minutes, verified by tocodynamometry) resulting in cervical dilatation and/or effacement.

For all patients, maternal age, gestational age, fetal and placental weights were recorded, and placental tissue was processed within 30 minutes of delivery. Placental villous tissue was immediately sampled from 5 random sites of the placenta and processed for trophoblast cell isolation as outlined below. Small villous pieces (∼5mm^3^) from all 5 sites were also flash frozen and stored at -80^0^C. Subsequently these villous tissue pieces from all 5 sites were powdered under liquid nitrogen and equal amounts from all sites were combined to be used for downstream analysis.

### Primary cell isolation and culture

The chorionic plate and decidua were removed from each of the randomly collected placental pieces leaving only villous tissue, which was thoroughly rinsed in PBS to remove excess blood. Primary cytotrophoblast were isolated from villous tissue using a protocol adapted from Eis *et al.* [35] using trypsin/DNAse digestion followed by density gradient purification. Isolated cytotrophoblast cells were then frozen in freezing media (10% DMSO in FBS) and stored in liquid nitrogen until usage. For culture, cytotrophoblast cells were rapidly thawed in a 37°C water bath and immediately diluted in Iscove’s modified Dulbecco’s medium (25mM glucose, 4mM glutamine and 1mM pyruvate) supplemented with 10% FBS and 1% penicillin/streptomycin (complete media). Cells were centrifuged at 1000g for 10 min and re-suspended in fresh complete media. Trophoblast cells were then plated in either 6 well-plates for collecting proteins and RNA or in 8 well chamber slides for staining. The cytotrophoblast (CT) when cultured differentiate into syncytiotrophoblast (ST) over a period of 72 hours. This differentiation is confirmed visually by observing cells under a light microscope. All studies were performed at two time points - 24 hrs denoted as CT stage and 72hrs – denoted as ST stage. Media was collected at both time points to measure secretion of βhCG as an additional way to confirm cytotrophoblast differentiation.

### Protein extraction and Western blotting

<u>Extraction from tissue:</u> Approximately 25mg of powdered tissue was mixed with 250μL of RIPA buffer supplemented with protease/phosphatase inhibitors (ThermoFisher Scientific, Cat. #A32959) and incubated on ice for 30 minutes. Post incubation, the tubes were centrifuged at 1000 rpm for 5 minutes and supernatant/lysate was transferred to a new tube. Total protein was quantified using the Pierce BCA Protein Assay Kit as per manufacturers protocol. (ThermoFisher Scientific, Cat. #23225).

<u>Extraction from cells</u>: Proteins were extracted from cultured trophoblast cells at 24 and 72hrs. Briefly, cells were washed in cold PBS with protease and phosphatase inhibitors (Halt^TM^ Protease Inhibitor Cocktail) followed by lysing in 150μL RIPA buffer supplemented with protease/phosphatase inhibitors (ThermoFisher Scientific, Cat. #A32959) for 20 minutes on ice. The lysate is then transferred to a 1.5ml tube and centrifuged at 1000g for 10 min at 4°C to remove cell debris. Total protein was quantified using the Pierce BCA Protein Assay Kit.

Approximately 30μg of protein was separated on 12% sodium dodecyl sulphate-polyacrylamide gel electrophoresis (SDS-PAGE) hand-cast gels for approximately 30 min at 30V followed by 2 hr at 100V and transferred for 1 hr at 100V onto nitrocellulose membranes using Mini-PROTEAN tetra cell electrophoresis chamber (BioRad, Cat. # 1658004). Membranes were blocked in 5% (w/v) nonfat milk in TBS + 0.1% Tween 20 (TBST) for 1 hr and incubated with primary antibodies (outlined in supplementary table 1) overnight at 4 °C. On the next day, membranes were washed three times in TBST for 5 min each and incubated with HRP-conjugated secondary antibodies. Membranes were washed and the antibody binding was detected using Supersignal West Pico Plus ECL Substrate (ThermoFisher Scientific, Cat. #34578) for 1 min and imaged using ChemiDoc Imaging System (Bio-Rad Laboratories, USA) and Image Lab V.5.1 software (Bio-Rad Laboratories, USA). Densities of immunoreactive bands were measured as arbitrary units by ImageJ software. Protein levels were normalized to a housekeeping protein β-actin (1: 20,000; Abcam, UK).

### Senescence associated β-galactosidase staining assay

Senescence-Associated β-Galactosidase (SA-βgal) is a lysosomal enzyme that cleaves β-D-galactose residues in β-D-galactosides. Detectable SA-βgal activity is the most extensively used marker for senescent or aging cells whether in culture or in mammalian tissues [36,37]. Senescent cells in trophoblasts isolated from LN and GDM placentae, CT were cultured for 24hrs in 8 chambered slides and then processed for SA-βgal staining using the CellEvent™ Senescence Green Detection Kit (Thermo Fisher) as per manufacturers protocol. Briefly, the cells were first fixed with 200uL of 1X fixative solution for 10 mins at RT after which the cells were washed with PBS containing 1% Bovine serum albumin. 200uL of prewarmed working solution was then applied to the cells. The slide chambers were sealed and incubated in dark for 2 hours in a CO_2_ free incubator at 37^0^C. After incubation, the working solution was drained, and cells were washed 3 times with 200 µL of PBS followed by staining with DAPI nuclear stain for 10 minutes. The cells were washed again once with PBS, the chamber walls separated, and the slides were coverslipped using SlowFade Diamond Antifade Mountant (Thermofisher Scientific, Cat. #S36972). After allowing to set, coverslips were sealed in place using clear nail polish. Images were captured using a Zeiss LSM 880 confocal microscope and processed using ImageJ Software. From each chamber, 3 randomly chosen areas were imaged. For each area, the total number of cells and cells positive for SA-βgal staining were counted to determine the % of senescent cells per sample.

### Enzyme linked Immunosorbent (ELISA) assay

The media collected from cultured trophoblast at 24 and 72hrs was assayed for levels of human chorionic gonadotropin β (βhCG) hormone using the Duo Set ELISA development kits (R&D Systems, USA) as per the manufacturers protocol. The optical density of the final-colored reaction product was measured at 450 nm using multispectral UV/VIS plate reader (Bio-Tek, VT). Standard curves were used to calculate protein content in the samples. The level of proteins detected was normalized to cellular protein.

#### Statistical analysis

The data was analyzed for statistical difference using the GraphPad Prism 7.0 software. All analysis consisted of n=5 samples per sex per clinical group. Outliers were detected using the ROUT method and wherever detected maximum 1 outlier was excluded from analysis. The values between CT and respective ST were analyzed using paired t-test whereas differences between groups were analyzed using ANOVA followed by post hoc Tukey analysis.

## RESULTS

### I. Clinical characteristics

The clinical characteristics of patients for which placental tissue and isolated trophoblast were utilized in the study are outlined in Table 1.0. There was no difference in maternal age and gestational age between the normoglycemic (NW) and type A2 GDM (GDM) group. However, patients from the GDM group had significantly higher BMI. As obesity increases the incidence of GDM by 3 times, we expected to see this in our GDM cohort [8,38,39]. All neonates had good outcomes and there was no difference in placental or fetal weight at birth between the NW and GDM groups.

**Table 1:** Clinical characteristics of study participants. Data presented as mean ± SD. Significant differences between NW and GDM groups were determined using the student’s *t* test. * *p* < 0.05 for NW vs. GDM, ^ϯ^ p<0.05 for same sex comparison between the NW and GDM groups.

|  | Normoglycemic<br>(NW, n=10) |  | Gestational Diabetes<br>(GDM, n=10) |  |
| --- | --- | --- | --- | --- |
|  | Male<br>(n=5) | Female<br>(n=5) | Male<br>(n=5) | Female<br>(n=5) |
| **Maternal age** (yrs) | 33.8 ± 3.5 |  | 34.7 ± 4.5 |  |
|  | 35 ± 2.2 | 32.6 ± 4.5 | 36 ± 3.7 | 33.4 ± 5.3 |
| **Maternal BMI**<br>(wt[kg]/height[m<sup>2</sup>]) | 21.29 ± 1.8 |  | 34.77 ± 4.8* |  |
|  | 22.5 ± 1.4 | 20.0 ± 1.2 | 35.2 ± 3.0*,† | 34.2 ± 6.5*,† |
| **Gestational age**<br>(wks) | 38.62 ± 0.90 |  | 38.08 ± 1.15 |  |
|  | 38.84 ± 0.93 | 38.4 ± 0.92 | 38.32 ± 0.93 | 37.84 ± 1.36 |
| **Fetal birthweight**<br>(g) | 3459 ± 319 |  | 3212 ± 258 |  |
|  | 3556 ± 214 | 3361 ± 399 | 3326 ± 205 | 3098 ± 275 |
| **Placental weight**<br>(g) | 556.4 ± 107.7 |  | 638.0 ± 150.5 |  |
|  | 609.7 ± 79.1 | 503.1 ± 112.9 | 690.8 ± 174.3 | 572.0 ± 97.3 |
| **Fetal:Placental**<br>**weight ratio** | 6.3 ± 1.1 |  | 5.5 ± 1.3 |  |
|  | 5.8 ± 0.6 | 6.8 ± 1.3 | 5.5 ± 1.6 | 5.6 ± 1.0 |

### II. GDM placentas have elevated levels of senescence marker p16

Placental villous tissue from NW and GDM women was used to determine the expression of senescence markers p21^WAF1/CIP1^ (p21) and p16^INK4A^(p16). We observed that expression of p21 was similar between the NW and GDM placentas whereas expression of p16 was significantly higher in GDM than NW (Fig 1A).

**Figure 1:**
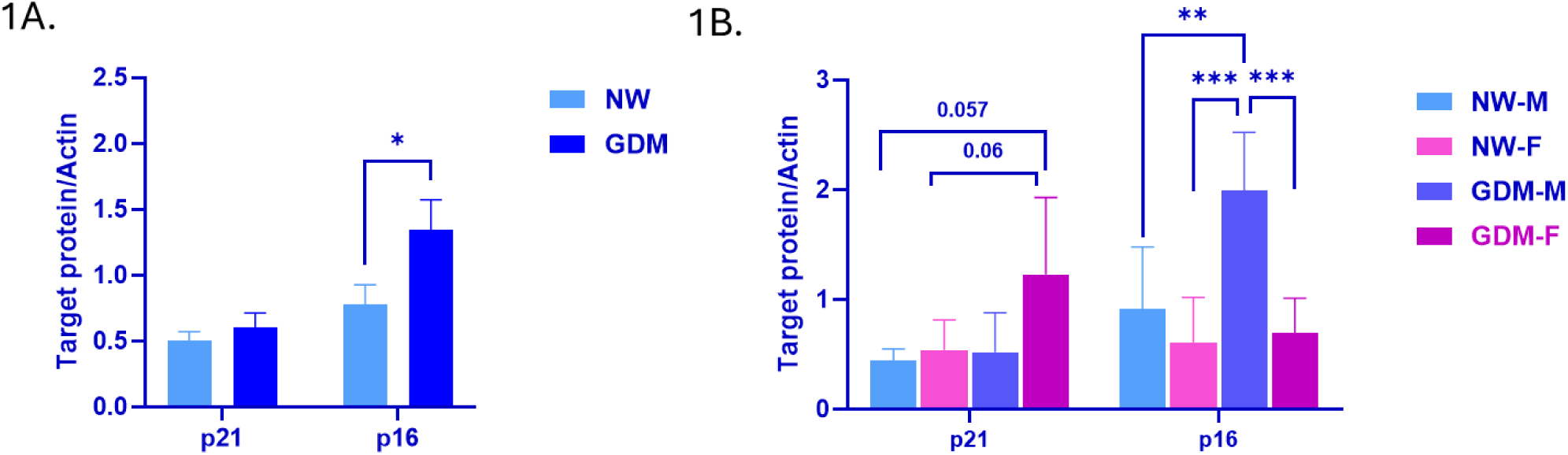
Protein expression of senescence markers p21 and p16 in villous tissue. **A.** Expression in NW and GDM villous tissue **B.** Expression in NW and GDM tissues stratified by fetal sex. Data presented Mean ± SEM with N=10 per clinical group with each containing n=5 males and n=5 females. * p<0.05, **p<0.01, *** p<0.001.

The data was then stratified based on fetal sex creating 4 subgroups: NW-male, NW-female, GDM-male and GDM-female. This analysis showed that expression of p21 was higher in the GDM-female compared to the other 3 subgroups, whereas the expression of p16 in the GDM-male subgroup was significantly higher compared to the other 3 subgroups. (Fig 1B). As outlined earlier, the women from the GDM group had significantly higher BMI compared to NW. We therefore also assessed expression of p21 and p16 in placentas from obese women with no type A2GDM to delineate if the changes seen in GDM placentas were due to maternal obesity. Interestingly obese placentas showed a pattern similar to the NW group (Supplementary figure 1) implying the changes in GDM placentas were specific to maternal diabetes. For the remaining experiments, we therefore continued using the NW group as our control.

### III. Trophoblast isolated from GDM placentae have higher numbers of senescent cells

As trophoblast are the major cell type that display characteristics of senescent cells in the placenta, we focused on them [40]. Cytotrophoblast isolated from NW and GDM placentas were cultured for 24 hours and the number of SA-β-gal positive cells was assessed at CT stage (Supplementary figure 2 shows representative images). We observed that CT from GDM placentas had a higher percentage of SA-β-gal positive cells compared to NW (Fig 2A). Stratifying this data by fetal sex showed that NW-male and GDM-male subgroups had comparable number of senescent cells, but the GDM-female subgroup had a higher number of senescent cells compared to the other 3 subgroups (Fig 2B).

**Figure 2:**
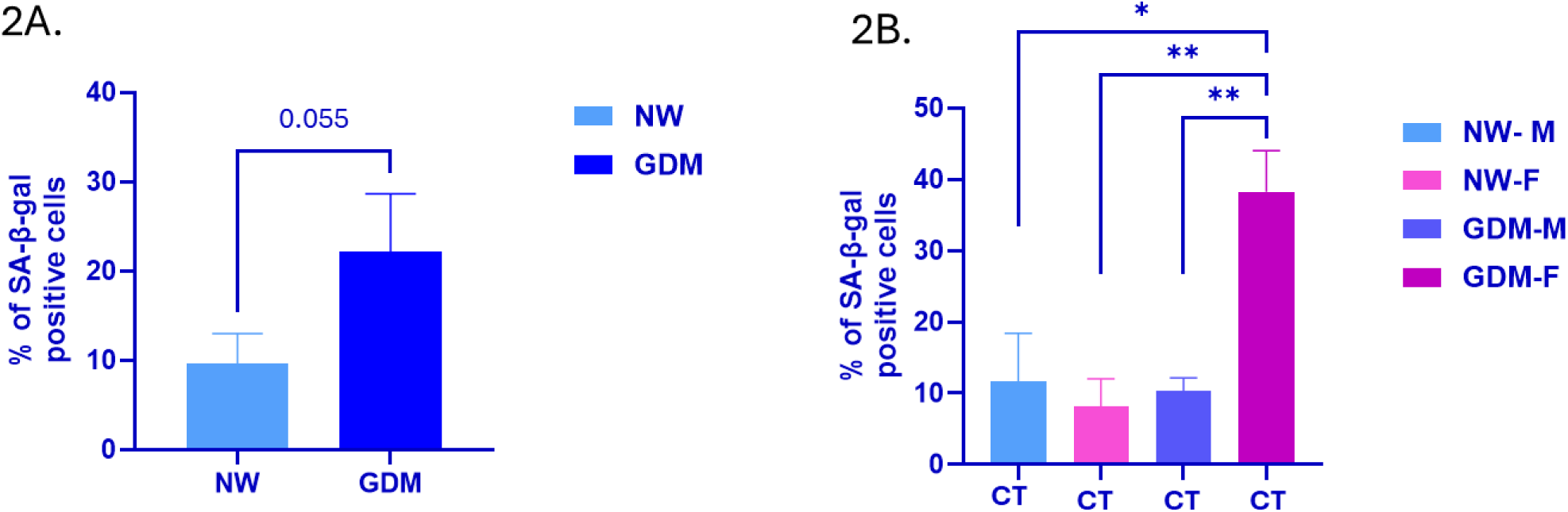
Percentage of senescent (SA-β-gal positive) cytotrophoblast after 24 hours of culture. **A.** Percentage of SA-β-Gal positive cells in CT isolated and cultured from NW and GDM placenta **B.** Percentage of SA-β-Gal positive cells stratified by fetal sex. N=6 per clinical group with each containing n=3 males and n=3 females. * p<0.05, **p<0.01 between groups

### IV. Differentiating cytotrophoblast from GDM placentas have distinct expression of senescence markers

We assessed the expression of p16 and p21 in cultured trophoblast at CT stage and post differentiation at ST stage. We observed that the levels of both p21 and p16 were lower in ST when compared to their respective CT in both NW and GDM groups - a trend previously identified in cells undergoing senescence [41] (Fig 3A, C). Comparison between the groups showed that CT from GDM group had significantly lower expression of p21 and higher expression of p16 (Fig 3A, C) compared to CT from NW group implying that trophoblast from both groups were undergoing senescence during trophoblast differentiation, but via activation of different pathways.

**Figure 3:**
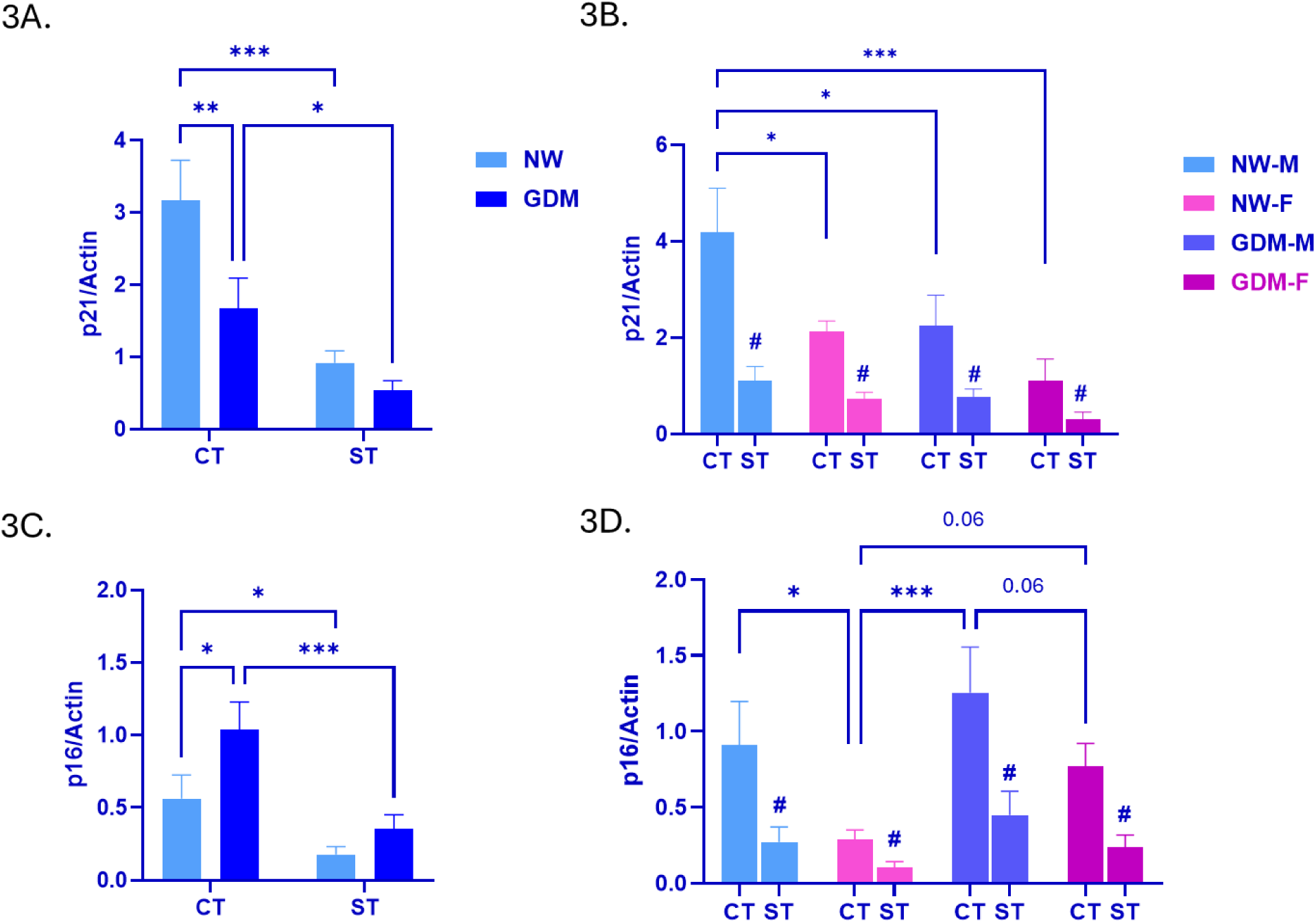
Protein expression of senescence markers p21 and p16 in CT and differentiated ST. Expression of p21 in NW and GDM CT and ST in **A**. sex combined manner, **B.** fetal sex stratified manner. Expression of p16 in NW and GDM CT and ST in **C**. sex combined manner, **D.** fetal sex stratified manner. Data presented Mean ± SEM with N=10 per clinical group with each containing n=5 males and n=5 females. * p<0.05, **p<0.01, *** p<0.001 between groups and ^#^ p<0.05 for comparison between CT vs ST in the same group.

Fetal sex-stratified analysis showed that the expression of p21 significantly declined in ST compared to their respective CT in all 4 subgroups (NW-male, NW-female and GDM-male, GDM-female). CT from the NW-male subgroup had higher p21 levels compared to NW-female as well as GDM-male and GDM-female subgroups (Fig 3B). Expression of p16 followed a similar trend wherein its levels also significantly declined in ST compared to their respective CT in all subgroups. NW-female CT also had significantly lower expression of p16 compared to CT from NW-male and GDM-male subgroups (Fig 3D).

### V. The p53-p21 and p16-pRB pathways are differentially activated in GDM trophoblast

To understand the mechanism behind differential senescence pathway activation, we assessed the expression of proteins, Tumor protein 53 (p53) and Retinoblastoma (RB) which act upstream and downstream of p16 and p21 respectively [42,43]. Phosphorylation of p53 is needed for its activation, whereas RB needs to be hypo-phosphorylated for mediating cell cycle arrest. We therefore compared the ratio of phosphorylated forms to total protein levels to determine their levels of activation.

We observed a higher proportion of activated p53 in GDM compared to NW at both the CT and ST stage (Fig 4A) implying higher activity of p53 in GDM trophoblast. Levels of phosphorylated 3 (normalized to Actin) were higher in ST from GDM compared to ST from NW group respectively (Supplementary Fig 3A). Similarly, total protein levels of p53 (normalized to Actin) were also significantly higher in ST from GDM compared to NW, whereas their levels in CT from both groups remained comparable between (Supplementary Fig 3C). Fetal sex-stratified analysis showed that the proportion of activated p53 declined in ST compared to their respective CT in NW-male and GDM-male subgroups but the proportions in NW-female and GDM-female subgroups were comparable (Fig 4B). Levels of phosphorylated p53 (normalized to Actin) were higher in ST from GDM-female compared to ST from the other 3 subgroups (Supplementary Fig 3B). Total protein levels of p53 were higher in ST from GDM-male and female groups compared to their respective CT, but this was not observed in NW-male or NW-female subgroups (Supplementary Fig 3D). The proportion of phosphorylated RB increased in ST from both groups compared to their respective CT (Fig 4C) however, GDM ST had significantly lower levels compared to NW ST, suggesting hypo-phosphorylation of RB in GDM trophoblast. The levels of phosphorylated RB (normalized to Actin) were lower in CT of GDM group compared to NW, but the levels in ST were comparable (Supplementary Fig 4A). The levels of total RB protein (normalized to Actin) showed a similar trend, with ST from both groups having lower levels compared to their respective CT (Supplementary Fig 4C). Fetal sex-stratified analysis showed that both male and female ST from NW group have a significantly higher proportion of phosphorylated RB compared to their respective CT and ST from NW-male subgroup had significantly higher proportion compared to ST in the other 3 subgroups (Fig 4D). These results suggest that during trophoblast differentiation, NW increased phosphorylation of Rb potentially to reduce its activity which was not attained in GDM. Levels of phosphorylated RB (normalized to Actin) were comparable in CT and ST in the subgroups except in NW-female wherein ST had significantly lower levels compared to their CT (Supplementary Fig 4B). For total RB protein (normalized to Actin), CT and ST from GDM-male subgroup had comparable levels whereas the remaining 3 subgroups had significantly lower levels in ST compared to their respective CT (Sup Fig 4D).

**Figure 4:**
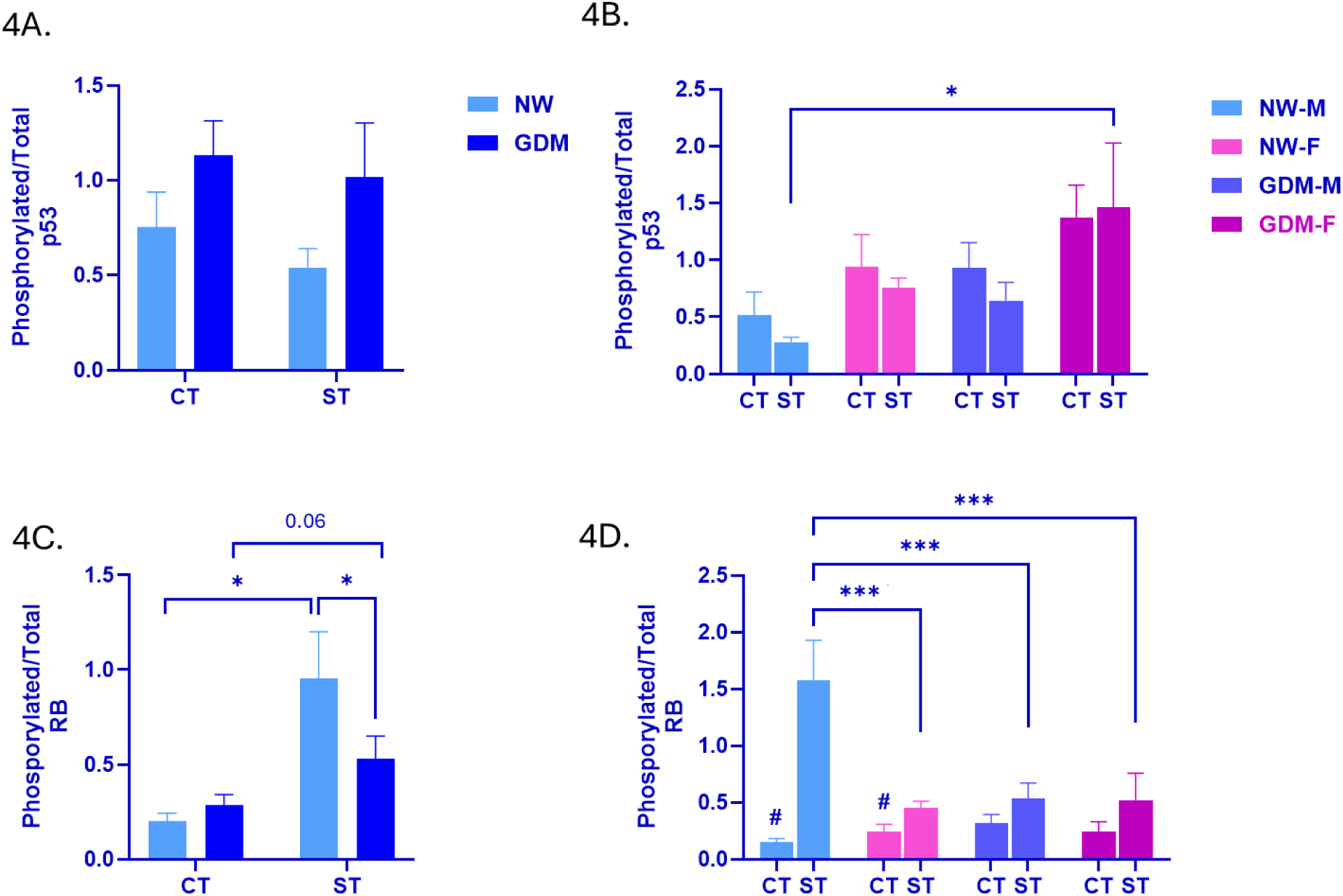
Protein expression of senescence pathway mediators p53 and RB in CT and differentiated ST. Ratio of expression of phosphorylated p53/total p53 in NW and GDM CT and ST in **A**. sex combined manner, **B.** fetal sex stratified manner. Ratio of expression of phosphorylated RB/total RB in NW and GDM CT and ST in **C**. sex combined manner, **D.** fetal sex stratified manner. Data presented Mean ± SEM with N=10 per clinical group with each containing n=5 males and n=5 females. * p<0.05, **p<0.01, *** p<0.001 between groups and ^#^ p<0.05 for comparison between CT vs ST in the same group.

### VI. Expression of Lamin A/C is high in GDM trophoblast

Lamins are a type of intermediate filament protein that line the inner surface of the nuclear envelope and contribute to the size, shape, and stability of the nucleus which is critical in multinucleated cells like the syncytiotrophoblast [44]. However, elevated levels of Lamin A/C were shown to mediate p53 induced senescence via destabilizing nuclear structure [45,46]. We observed significantly higher expression of Lamin A/C in both CT and ST from GDM placentas compared to CT and ST from NW placentas respectively (Fig 5A)). Fetal sex stratified analysis showed that CT and ST in both NW-male and NW-female subgroups had comparable expression. Expression in CT from GDM-female subgroup was significantly higher compared to CT from NW-male and NW-female subgroup whereas ST from GDM-male subgroup had significantly higher expression compared to ST from the NW-male and NW-female subgroup (Fig 5B).

**Figure 5:**
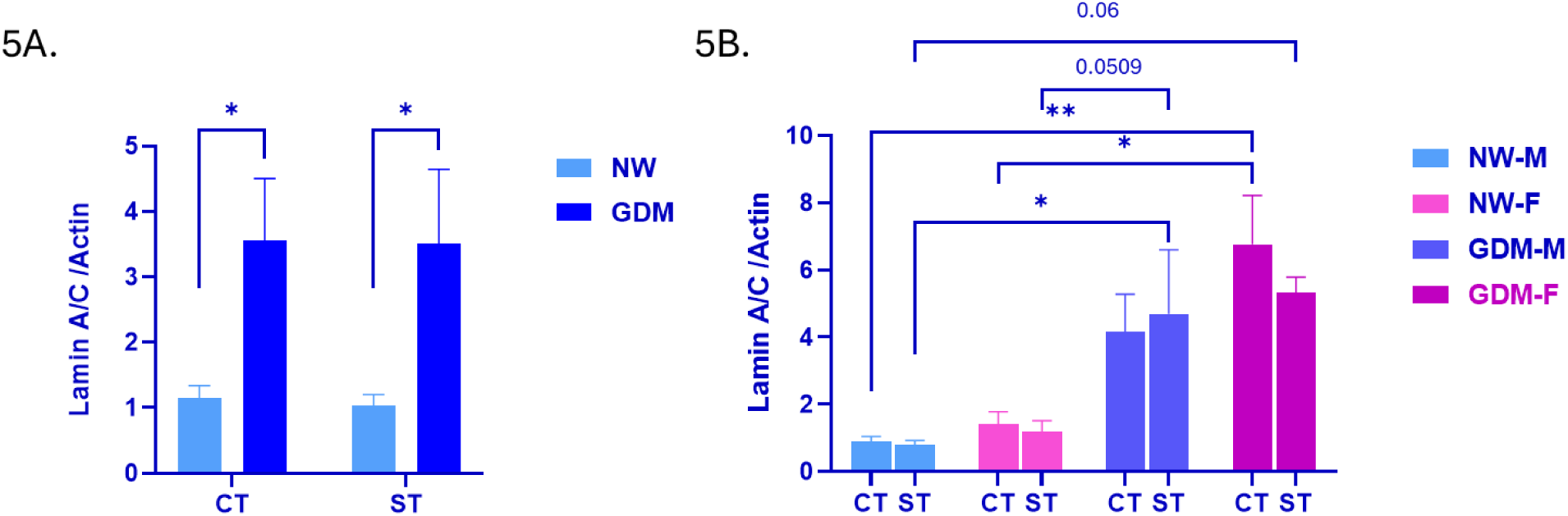
Protein expression of Lamin A/C in CT and differentiated ST. Expression of Lamin A/C in CT and ST **A**. sex combined manner, **B.** fetal sex stratified manner. Data presented Mean ± SEM with N=10 per clinical group with each containing n=5 males and n=5 females. * p<0.05, **p<0.01, *** p<0.001 between groups.

### VII. Expression of both GCM1 and βhCG is reduced in differentiating trophoblast from GDM placentas

A recent report suggested that p21 is involved in regulation of Glial cell missing -1 (GCM1) implying a physiological link between activation of senescence and trophoblast differentiation [47]. We observed that levels of GCM1 were significantly lower in ST from both groups compared to their respective CT suggesting induction of trophoblast differentiation in both groups. However, both CT and ST from GDM placentas had significantly lower levels of GCM1 compared to CT and

As GCM1 is known to regulate human chorionic gonadotropin-β (βhCG) – also involved in trophoblast differentiation, we next evaluated βhCG secretion from these cells. We observed that secretion of βhCG significantly increased as CT differentiated to ST in trophoblast from NW and GDM placentas, in line with previous literature and again implying induction of differentiation in CT from both groups (Fig 6C). However, the βhCG levels secreted GDM ST by them were significantly lower than ST from NW placentas implying reduced trophoblast differentiation. Fetal sex stratified analysis showed that βhCG secretion significantly increased in ST compared to their respective CT in all four subgroups. Interestingly, ST from the NW-female subgroup had significantly higher levels compared to ST from NW-male but the levels secreted by ST from GDM-males and GDM-female subgroups were comparable suggesting a sexual dimorphic pattern in normoglycemic placentas which was absent in GDM. The **β**hCG levels secreted by ST from the NW-female subgroup were also significantly higher than GDM-male and female subgroups (Fig 6D).

**Figure 6:**
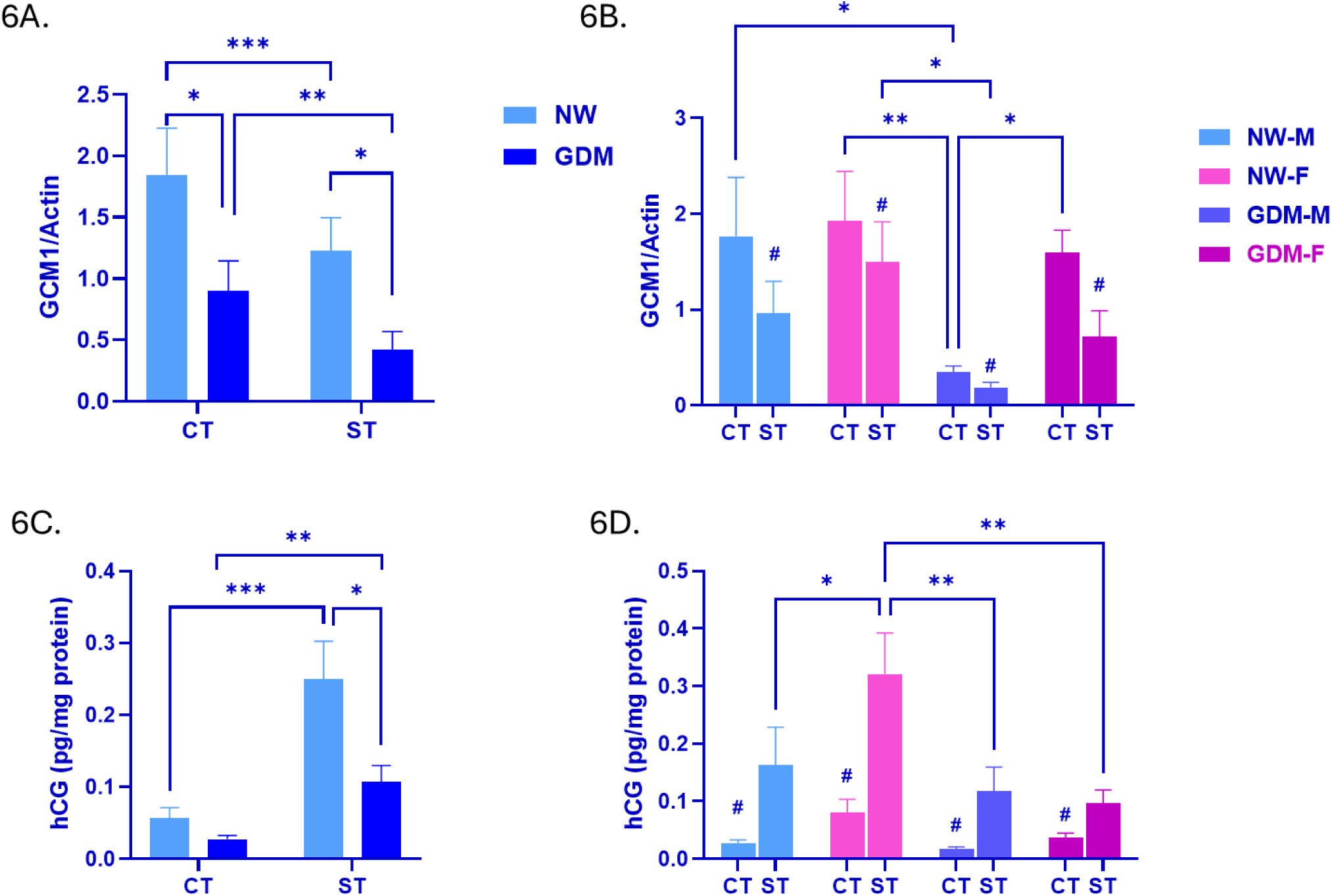
Protein expression of GCM1 and βhCG secretion in CT and differentiated ST. Expression of GCM1 CT and ST from NW and GDM **A**. sex combined manner, **B.** fetal sex stratified manner. Levels of βhCG secreted in media by CT and ST from NW and GDM **C**. sex combined manner, D. fetal sex stratified manner Data presented Mean ± SEM with N=10 per clinical group with each containing n=5 males and n=5 females. * p<0.05, **p<0.01, *** p<0.001 between groups and # p<0.05 for comparison between CT vs ST in the same group.

## DISCUSSION

The current literature exploring senescence in human placenta focuses mainly on expression of senescence associated markers with several studies reporting upregulation in post mature placentas, and in placentas from pregnancies complicated by PE, FGR and stillbirth [31,32,48]. However, the functional role of senescence in placental tissue remains unexplored. Here, using the primary trophoblast cell model we demonstrate that while senescence pathways are induced during physiological trophoblast differentiation, their induction is reduced in the maternal condition of GDM and affects differentiation which in turn might affect placental function. We also assessed sexual dimorphism in these pathways which has not been previously addressed.

Cellular senescence is often synonymously used with ageing as accumulation of senescent cells in tissues has been observed in aging and age-related pathologies [49]. However, senescence also plays crucial physiological roles in maintaining tissue homeostasis, wound healing and as an anti-cancer mechanism in tissues [50]. Cellular senescence has also emerged as a response to different stressors including genotoxic stress, nutrient imbalance and mitochondrial dysfunction. The latter two being extensively reported in GDM placentas [17,51]. Programmed cellular senescence was shown to be crucial for embryonic development and was also proposed to play a role in timing of parturition [31]. Studies exploring aging and/or senescence in placental tissues have reported an increase in p21, p16 and p53 markers in pregnancies complicated by PE, IUGR, stillbirth and miscarriage [31,32,48,52]. Increase in senescence markers was also reported to coincide with increasing oxidative stress in these pathological states as well as post mature placentas [32]. Most previous studies were conducted using villous tissues from these pregnancies or trophoblast cell lines of cancerous origins that might have already bypassed cell cycle-related processes which may be dysregulated in senescence *in-vivo* [31,32,48,40,53]. Biron-Shental *et al* reported an increase in the senescence markers – SA-β-gal staining, senescence-associated heterochromatin foci and reduced telomere lengths in GDM placentas [33]. Garcia-Martin *et al* also reported reduced telomere lengths in GDM placentas, which was reversed when mothers received interventions such as metformin or insulin but focused only on placentas from male fetuses [54]. While these studies support our findings, they lack the ability to distinguish between physiological and pathological senescence as they rely on histological markers or protein levels in bulk placental tissue and do not consider sexual dimorphism. Higuchi *et al* performed a longitudinal study across placental tissue from 1^st^, 2^nd^ and 3^rd^ trimester and reported higher senescence in CT from early to mid-pregnancy in association with syncytial fusion, while senescence was observed in ST in late pregnancy. The authors suggested this change in senescence across gestation may play a role in maintaining placental function [53], however the physiological role of senescence in placental function remains unevaluated in GDM. In our analysis with villous tissue, we also observed altered expression of senescence regulators p21 and p16 in GDM placentas. P21 and p16 are cyclin-dependent kinase (CDK) inhibitors that prevent cell cycle progression by interacting respectively with the transcription factors p53 and RB that regulate expression of genes involved in nutrient metabolism, DNA replication, cell growth and apoptosis (Fig. 7) [55]. We observed an increase in expression of p16 in GDM placentas compared to NW controls implying an increase in p16 mediated senescence in GDM. We also observed sexual dimorphism in expression, wherein male placentas from the GDM group had significantly higher levels of p16 compared to female placentas. These results suggest that male and female placentas differ in activation of senescence pathways in response to maternal GDM conditions. As trophoblast display characteristics of senescent cells, we focused on them and utilized cytotrophoblast isolated from pregnancies complicated by NW and GDM to study expression of senescence pathway mediators as they differentiated to syncytiotrophoblast [40]. This also enabled us to understand differences between physiological senescence in NW and how it is altered in GDM pathology. At 24 hours post culture, we observed SA-β-gal positive cells in both NW and GDM CT with GDM CT having a significantly higher number suggesting premature activation of senescence pathways. We also observed that CT from both groups activated both p16 and p21 mediated senescence as part of their differentiation process and their levels declined as CT differentiated to ST (Fig. 1), which is synonymous with observations made in non-placental senescent cells wherein levels of p21 and p16 decline once the senescence program is initiated [41]. P21 is proposed to be activated as part of the developmental senescence program as p21 mutant mice showed defects in embryonic senescence among other developmental defects which would explain the robust expression of p21 in CT from both sexes in NW and GDM placentas [19,56]. Taken together, these results further solidify the role for physiological senescence in trophoblast differentiation. Studies from other models have also suggested that p21 is activated in the initial stages of senescence whereas p16 is activated in the late stages and an increase in expression of p16 has been linked to a senescence-mediated decline in progenitor stem cell populations and function [57,58]. We observed lower p21 expression and higher p16 levels in GDM CT compared to CT from NW placentas. P16 also showed a sexually dimorphic pattern wherein female CT from both NW and GDM groups had lower expression compared to their respective male CT. Reduced levels of p21 and higher levels of p16 in CT from GDM trophoblast with higher number of SA-β-gal positive cells could indicate disruption in physiological senescence in these placentas.

**Figure 7:**
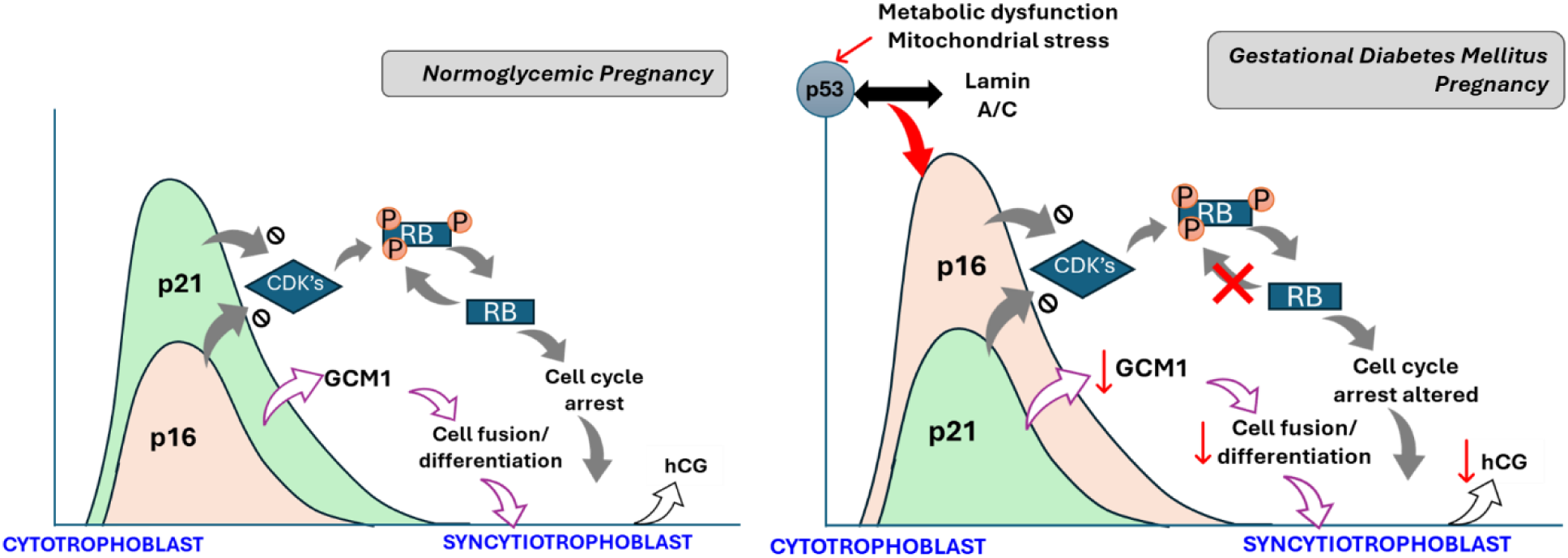
Overview of changes in senescence pathways as cytotrophoblast from NW and GDM placentas differentiate into syncytiotrophoblast.

We also observed an increase in activated p53 in which typically acts upstream of p21 in both GDM CT and ST. Cellular stressors including metabolic stress, oxidative stress and mitochondrial stress can all activate p53 as a way to combat that stress and can lead to cell death [59,60]. The non-significant increase in p53 observed here could be in response to cellular stress in GDM. However, increased activation of p53 did not coincide with a concomitant increase in p21 as expected, which might be because p21 can be regulated in a p53 independent manner [61,62]. However, p53 was recently shown to induce p16 expression via Lamin A/C stabilization [45]. Indeed, we observed increase in Lamin A/C levels in both CT and their differentiated ST from GDM placentas suggest that even though there was a nominal activation of p53, this was enough to induce p16. While there were no specific sexually dimorphic trends in expression of p53 and Lamin A/C, it is worth highlighting that both CT and ST from GDM females displayed higher expression of both proteins. We also assessed phosphorylated and total levels of RB protein which acts downstream of p16. P16 enforces senescence by inhibiting CDK4/6, leading to RB hypo-phosphorylation which results in its binding to E2F’s forming a repressive complex that binds and prevents expression of genes needed for cell cycle progression [63,64] (Fig 7). In our analysis we observed that CT from both NW and GDM groups had comparable proportion of phosphorylated/total RB which increased in their respective ST. This increase in phosphorylated/total RB could be explained by reduced levels of total RB protein implying rapid degradation of RB once the cell cycle arrest pathways have been activated. Interestingly, we observed that ST from the GDM group were more hypo-phosphorylated compared to ST from the NW group which aligns with increased cell cycle arrest and senescence in GDM ST [43].

Additionally, p21 is also known to regulate RB phosphorylation and formation of the repressive RB-E2F complexes. P21 mediated inhibition of CDK’s also contributes to reduced phosphorylation of RB and progress to cell cycle arrest [65]. The decline in p21 levels as CT from NW placentas differentiate to ST coincides with an increase in RB phosphorylation status in these cells highlighting the role of the p21-RB axis in physiological senescence. However, low levels of p21 in GDM placental CT with a further decline in ST suggest that this axis is also dysregulated in GDM trophoblast. To determine if this dysregulation in senescence affects placental function, we analyzed levels of trophoblast differentiation factors GCM and βhCG. Expression of GCM1 was significantly downregulated in both CT and ST from the GDM group compared to the CT and ST from NW group respectively, suggesting reduced differentiation. As CT from both groups differentiated to ST, they increased their secretion of βhCG confirming differentiation however, the βhCG levels secreted by GDM ST were significantly lower compared to ST from the NW group again confirming dysregulated differentiation in GDM trophoblast. GCM1 may induce p21 expression to arrest the cell cycle, and in turn p21 enhances the function of GCM1 to begin the fusion program [47]. The reduced expression of both p21 and GCM1, and the downstream target βhCG suggest that the dysregulation in p21 mediated physiological senescence in GDM placentas affects trophoblast differentiation. This dysregulated differentiation could explain the presence of immature villi observed in GDM placentas [13]. Taken together, our results suggest that GDM cytotrophoblast may be undergoing premature senescence, driven by p53 mediated Lamin A/C accumulation and p16/RB activation (Fig 7). Our results also suggest that loss in p21 in GDM trophoblast alters physiological senescence and is capable of disrupting trophoblast differentiation by altering GCM1 and βhCG levels. Disruption in trophoblast differentiation can alter placental function in numerous ways such as altered secretion of hormones and signaling mediators, altered expression of nutrient transporters, etc. This is the first study to outline disruption of placental senescence pathways in GDM and their potential impact on trophoblast differentiation. Several stressors including oxidative stress, mitochondrial dysfunction, and metabolic stress have all been shown to induce senescence pathways in non-pregnant individuals tissues [66,67,27,22–26]. GDM placentas must adapt to the altered maternal metabolic environment and have been reported to have increased oxidative stress as well as mitochondrial dysfunction [68–70]. It is plausible that a combination of these conditions could alter physiological senescence in GDM placentas (mediated via p21) and induce premature elevated senescence (mediated by p16). We also report sexual dimorphism in placenta senescence pathways further highlighting how fetal sex plays a role in shaping placental function and its response to maternal stress. Our study warrants further research into the potential stimulus for the reported dysregulated placental senescence in GDM and the sexual dimorphism therein.

## AUTHOR CONTRIBUTIONS

LK conceptualized, designed and performed the experiments. LK was also responsible for data generation and analysis. NM was involved with identifying patients and KC and KA were involved with sample collection and running experiments. LK wrote the manuscript with contribution from LM.

