## Supplementary for "Differential activation of p53-Lamin A/C and p16-RB mediated senescence pathways in trophoblast from pregnancies complicated by type A2 Gestational Diabetes Mellitus"

**Supplementary Table 1.0**

| Antibody | Host species, clonality | Company |
|---|---|---|
| Anti-GCM1 | Rabbit, polyclonal | AVIVA Biosystems |
| Anti-p21$^{Waf1/Cip1}$ | Rabbit, monoclonal | Cell signaling |
| Anti-p16 $^{INK4}$ | Rabbit, monoclonal | Cell signaling |
| Anti-phosphorylated p53 | Rabbit, polyclonal | Abcam |
| Anti-p53 | Rabbit, monoclonal | Cell signaling |
| Anti-phospho-Rb (Ser807/811) | Rabbit, monoclonal | Cell signaling |
| Anti-Rb | Mouse, monoclonal | Santacruz |
| Anti-Lamin A/C | Mouse, monoclonal | Santacruz |

**SUPPLEMENTARY MATERIAL:**

**Supplementary figure 1**. Expression of p21 and p16 in male and female placentas from normoglycemic women (NW), women who are obese only (OB) and women with type A2GDM (GDM). Data expressed as Mean ± SEM with n=5 per sex, per group.

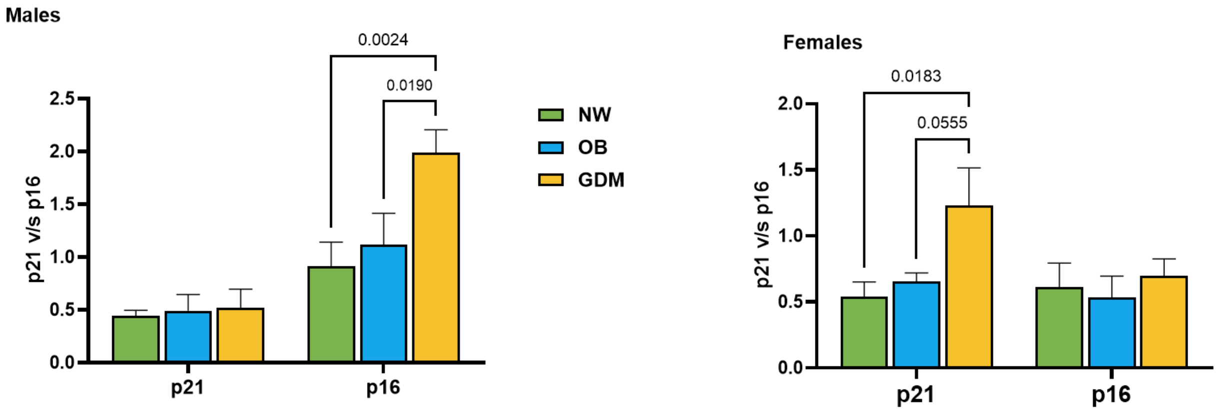

**Supplementary figure 2:** Representative images of SA-β-gal staining at **A**.10X and **B**. at 40X magnification. White arrows indicate SA-β-gal positive cells and white asterisks represent SA-β-gal negative cells.

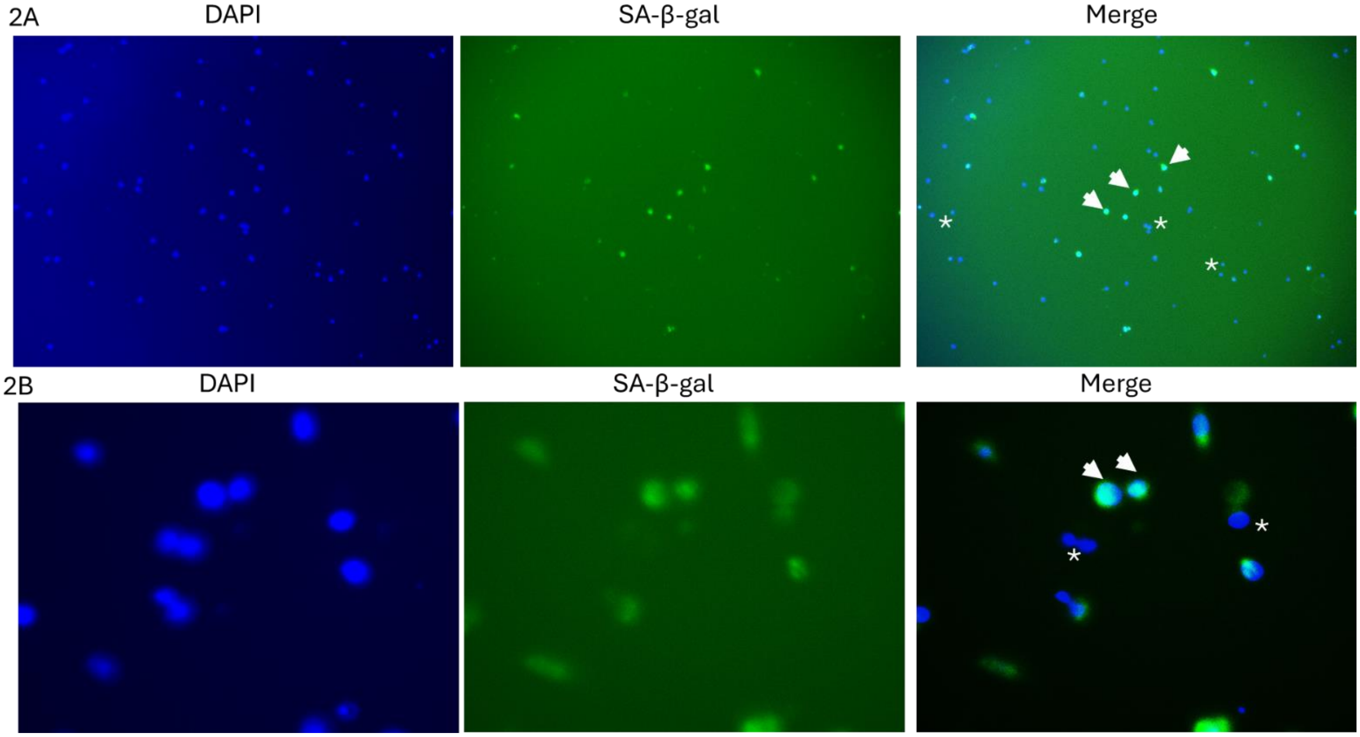

**Supplementary figure 3: A** Expression pattern of phosphorylated p53 (normalized to Actin) and Total p53 (normalized to Actin) in CT and ST from NW and GDM placenta respectively in **A,C**. sex combined manner, **B, D.** fetal sex stratified manner.

Data presented Mean ± SEM with N=10 per clinical group with each containing n=5 males and n=5 females. * p<0.05, **p<0.01 between groups and # p<0.05 for comparison between CT vs ST in the same group.

Supplementary figure 3

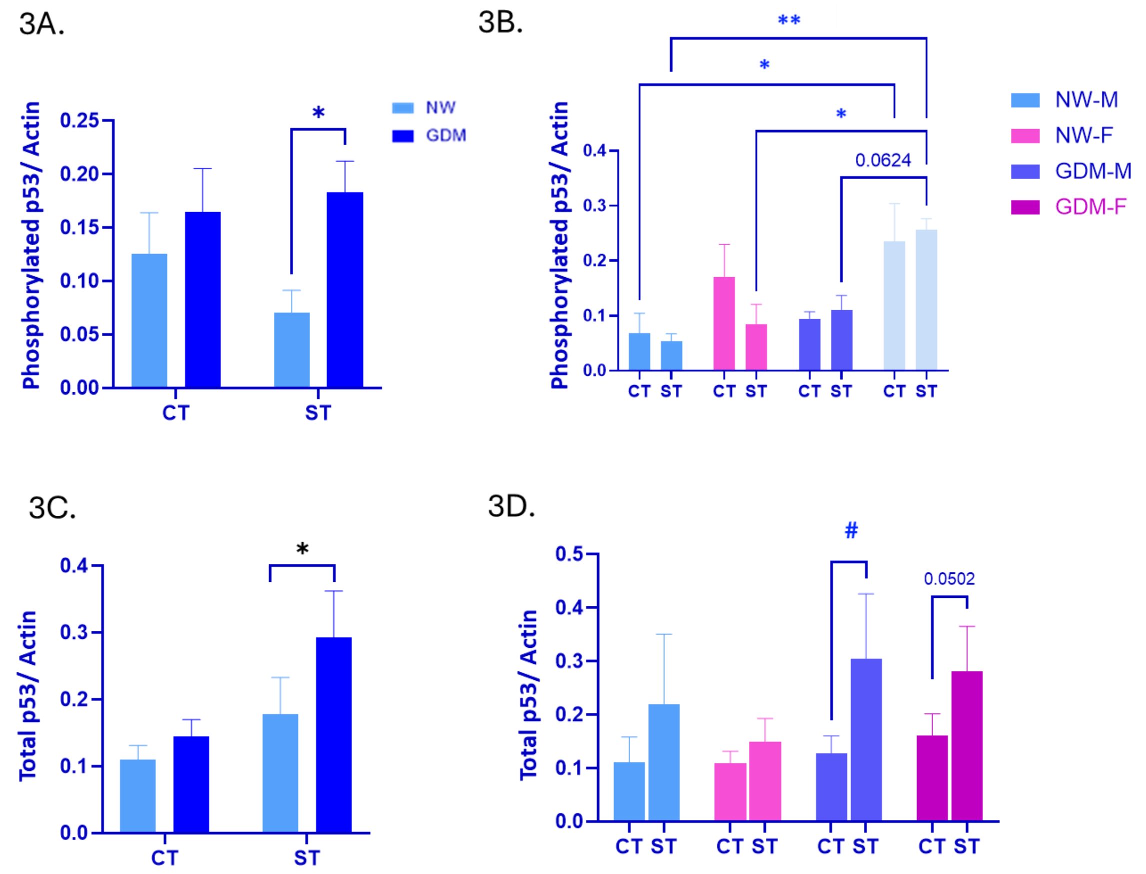

**Supplementary figure 4:** Expression pattern of phosphorylated RB (normalized to Actin) and Total RB (normalized to Actin) in CT and ST from NW and GDM placentas respectively in **A,C**. sex combined manner, **B, D.** fetal sex stratified manner.

Data presented Mean ± SEM with N=10 per clinical group with each containing n=5 males and n=5 females. * p<0.05 between groups and # p<0.05 for comparison between CT vs ST in the same group.

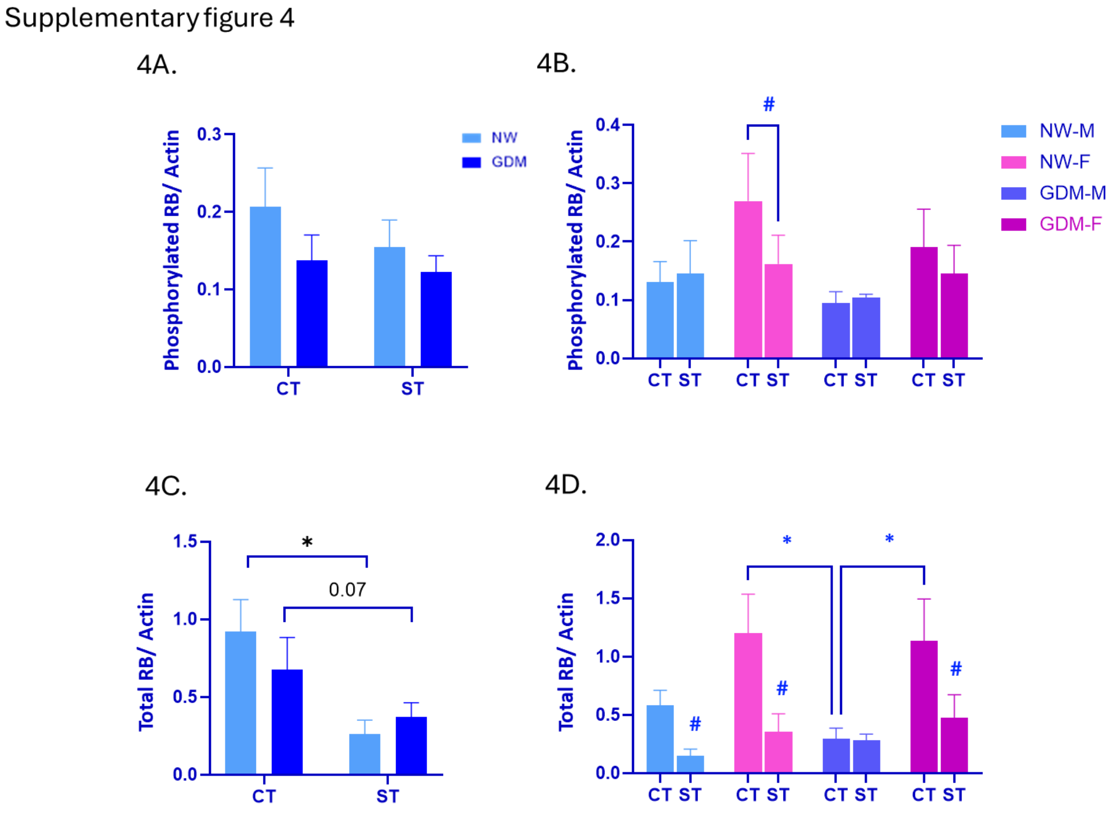
